# A heterologous expression system allows rapid testing and discovery of resistance-conferring mutations from multiple pathogenic fungi

**DOI:** 10.64898/2026.09.18.751866

**Authors:** David F. Jordan, Mathilde Roland-Caverivière, Camille Bédard, Alexandre K. Dubé, Isabelle Gagnon-Arsenault, Christian R. Landry

## Abstract

To face the fungal infections that affect millions of people every year, a better understanding of resistance-conferring mutations is necessary. One obstacle is the considerable time and effort necessary to construct point mutations in pathogenic species, which is required to validate their role in resistance. To overcome this problem, we have constructed a series of heterologous expression plasmids that allow expression of the *ERG3, ERG6* and *ERG11* orthologs from *Candida albicans, Candidozyma auris, Nakaseomyces glabratus, Candida parapsilosis, Candida tropicalis,* and the *CYP51A* and *CYP51B* genes of *Aspergillus fumigatus*. All orthologs are codon-optimized and synthesized for expression in the budding yeast *Saccharomyces cerevisiae*. We also constructed host strains in which the native *ERG* genes are either deleted or under the control of a repressible promoter. We showed that all orthologs complement the function of the native genes when expressed from the *S. cerevisiae* native promoters. We recreated previously reported resistance-conferring mutations, and showed that expression in *S. cerevisiae* replicates known resistance phenotypes. We then used the system to test corresponding mutations across orthologs, and showed that most resistance phenotypes are conserved. The heterologous expression system allows rapid construction of mutants, by taking advantage of the abundant genetic tools available for *S. cerevisiae.* Screening of mutants can be done within a few experimental steps, greatly increasing the throughput compared to the introduction of mutations in pathogenic fungi. As a proof-of-concept, we used the system to discover new resistance-conferring mutations, using random mutagenesis. All plasmids are made publicly available.

**Summary:** One problem impeding research into fungal pathogens is that putative resistance-conferring mutations are difficult to recreate and study. To facilitate research, we have built a series of heterologous expression plasmids for *ERG3, ERG6* and *ERG11* in multiple clinically relevant fungal species. We test the expression plasmids by recreating known phenotypes of resistance-conferring mutations. As a proof-of-concept, we use the system to discover new resistance-conferring mutations, notably using a random-mutagenesis method. Researchers working on fungal pathogens can now use this system to quickly and efficiently phenotype large numbers of mutations, allowing larger-scale studies than previously.

## Introduction

Yeasts and filamentous fungi related infections exact a massive toll on global human health (Case et al. 2025). While breakthroughs have taken place in the past few years, much more effort is necessary in order to fully understand the biology of pathogenic yeasts. Further exacerbating the problem is the emergence of pathogenic fungi which have become resistant to one or more of the commonly used antifungal drugs (Robbins et al. 2017). In fact, antifungal resistance is considered an emerging crisis, which can affect human life, animal life (both wild and livestock) and agricultural production (Fisher et al. 2018; Fisher et al. 2022). As only 4 classes of antifungal drugs are commonly used by physicians to treat invasive fungal infections, resistance to any drug drastically reduces treatment options (Susan et al. 2025). Resistance to antifungal drugs can occur through many mechanisms (Lee et al. 2023), but clinical and laboratory data suggests that it emerges mainly through point mutations in key genes (Jay et al. 2025).

Despite the importance of fungal infections and the prevalence of resistance-conferring mutations, these reported mutations are often relatively poorly characterized, often being simply reported from sequencing of clinical or laboratory samples without functional validation (Bédard et al. 2025). More thorough characterization of mutations of interest is hampered by the relative difficulty of laboratory work with fungal pathogens. While great strides have been made in the past decades, work on many pathogenic species is generally more time and resource intensive than work on model organisms, such as budding yeast (Ross and Santiago-Tirado 2024). Furthermore, some pathogenic yeast and other ascomycetes do not naturally contain plasmids, rendering persistence of plasmids rare (Ross and Santiago-Tirado 2024). Other difficulties include the absence or extreme rareness of a mating cycle, which precludes approaches based on mating and meiosis to recombine alleles of interest, and the fact that many pathogens are diploid, requiring more extensive genetic changes to ensure a homozygous genotype (Uthayakumar et al. 2020). Moreover, the pathogenic nature of the biological material requires biosafety precautions which can slow down research. Limited genetic engineering tools do exist that can produce genetic changes and point mutations in many species. For example, CRISPR-based approaches have been developed for many pathogenic yeast species (Maroc and Fairhead 2019; Uthayakumar et al. 2020; Santana and O’Meara 2021; Hartuis et al. 2024; Doorley et al. 2025). However, the available tools remain limited compared to the resources available for genetic engineering in model organisms such as *Saccharomyces cerevisiae*.

As shown by a recent systematic survey of resistance mutations (FungAMR, (Bédard et al. 2025)), one powerful approach to overcome genetic engineering limitations is to use heterologous expression in a model system. The FungAMR database lists 431 entries from 37 studies where antifungal resistance or susceptibility was tested using heterologous expression of a mutated gene (Bédard et al. 2025). Heterologous expression has proven useful in the past. As far back as 30 years ago researchers expressed the *CDR1* ortholog from *Candida albicans* in a modified *S. cerevisiae* strain to characterize this gene (Prasad et al. 1995; Nakamura et al. 2001). One specific example which has been implemented in multiple forms is expressing different alleles of *C. albicans ERG11* on a plasmid in *S. cerevisiae*, to more easily measure antifungal resistance (Sanglard et al. 1998; Chau et al. 2004; Xiang et al. 2013; Healey et al. 2018). Heterologous expression can also be achieved between orthologs of more distantly related species. For example, *Aspergillus fumigatus CYP51A* and *CYP51B* were used to complement the function of the native *S. cerevisiae ERG11*. This complementation also allowed the recreation of multiple point mutations, allowing measurement of antifungal resistance (Martel et al. 2010; Alcazar-Fuoli et al. 2011). Heterologous expression has also been used with the glucan synthase gene *FKS1*, where mutations are often clustered in well-defined “hotspots”, by replacing only the *S. cerevisiae FKS1* hotspot 1 with the orthologous hotspots from species across the ascomycete clade (Katiyar and Edlind 2009; Dudiuk et al. 2017; Durand et al. 2026). While heterologous expression is mostly undertaken in *S. cerevisiae*, this is not always the case, as in one study which expressed 9 different *ERG3* orthologs in *C. albicans* (Luna-Tapia et al. 2021). Pushing the approach even further, two recent studies have used heterologous expression in *S. cerevisiae* to test the drug resistance of libraries of thousands of mutant sequences of either *Pneumocystis jirovecii DHFR* (Rouleau et al. 2024) or *C. albicans ERG11* (Bédard et al. 2024).

As mentioned, many studies using heterologous expression systems have examined different orthologs of the *ERG11* gene. *ERG11* is a key player in the sterol synthesis pathway, which principally produces ergosterol, the main sterol component in fungal cellular membranes (Zinser et al. 1991). Different sterol synthesis genes play a role in antifungal resistance in multiple fungal species (Alcazar-Fuoli and Mellado 2012; Whaley et al. 2016; Ahmad et al. 2019; Asadzadeh et al. 2023). In particular, two of these, *ERG3* and *ERG6*, are involved in resistance not only to azole antifungals, but also to polyene and echinocandin class drugs (Gerstein et al. 2012; Bhattacharya et al. 2018; Spettel et al. 2019; Carolus, Sofras, Boccarella, Sephton-Clark, et al. 2024; Carolus, Sofras, Boccarella, Jacobs, et al. 2024). Resistance to azoles can occur through loss-of-function mutations in *ERG3*, though the FungAMR database reveals no such clear pattern for *ERG6* (Bédard et al. 2025). Mutations in *ERG3* have species-specific effects in conferring resistance to polyene drugs through loss-of-function mutations, and the FungAMR database reports echinocandin resistant yeast strains that contain loss-of-function mutations in *ERG3* (Bédard et al. 2025). Additionally, loss-of-function mutations in *ERG6* are known to confer resistance to both polyene drugs: amphotericin B and nystatin (Jensen-Pergakes et al. 1998; Vincent et al. 2013). The precise cellular mechanisms behind *ERG3* and *ERG6*’s involvement in echinocandin and polyene resistance is still unclear, though both genes play a role in the synthesis of ergosterol (Parks et al. 1995; Carolus et al. 2025). *ERG3* and *ERG6* loss-of-function mutations influence the proportion of ergosterol and other sterols present in the cell membrane (Geber et al. 1995; Jensen-Pergakes et al. 1998), and may in this manner play a role in antifungal resistance and susceptibility. On the other hand, for *ERG11* a much larger number of mutations are reported in the FungAMR database and the mechanisms of resistance are much better understood. This is not surprising, as *ERG11* is the most frequently mutated gene in cases of resistance to azole class antifungals (Bédard et al. 2025). Erg11 is the binding target of azoles, consequently, resistance-conferring mutations in *ERG11* weaken the binding of azoles to the Erg11 active site (Lupetti et al. 2002; Bédard et al. 2024). However, the antifungal action of azoles does not merely result from the inhibition of Erg11, but also by the resulting production of a toxic sterol through an alternative metabolic pathway involving both Erg3 and Erg6 (Eliaš et al. 2024). Still, the vast majority of *ERG11* mutants in the FungAMR database are characterized only for the *C. albicans* ortholog (Bédard et al. 2025). Further characterization of resistant *ERG3, ERG6* and different orthologs of *ERG11* could greatly increase our understanding of antifungal resistance as well as sterol synthesis.

Heterologous gene expression has the potential to be extremely powerful, and accelerate the screening of potentially resistance-conferring mutations in sterol synthesis pathway genes. However, previous efforts have all been piecemeal and each study has used their own specially designed strategy. To make heterologous expression more readily available to more researchers, and to standardize the approach, a systematized series of plasmids and yeast strains for heterologous expression of *ERG3, ERG6* and *ERG11* orthologs was constructed. For each of the three genes, plasmids expressing orthologs from major fungal pathogens *C. albicans, Candidozyma auris, Nakaseomyces glabratus, Candida parapsilosis, Candida tropicalis* as well as *A. fumigatus CYP51A* and *CYP51B*, were constructed, along with the *S. cerevisiae* ortholog as a control. Along with the series of plasmids, *ERG3* and *ERG6* deletion strains of *S. cerevisiae* have been constructed, as well as strains in which expression of the native *ERG3, ERG6* or *ERG11* gene can be repressed, allowing unbiased evaluation of antifungal resistance or susceptibility. Gene sequences were codon-optimized for expression in *S. cerevisiae* and their expression is driven by the native *S. cerevisiae* promoter and terminator. The plasmids and strains are available to the research community. Here we describe the construction of the aforementioned plasmids and strains. We then show that all orthologs complement the *S. cerevisiae* gene, and the phenotypic effects of resistance from pathogenic species is measured by recreating resistance-conferring mutations on the heterologous expression plasmids. The system is then used to rapidly evaluate the antifungal resistance of uncharacterized mutations across all orthologs. Finally, we demonstrate a further use of heterologous expression by undertaking a random mutagenesis strategy to discover novel resistance-conferring mutations in *C. albicans ERG11*.

## Results

### Multiple *ERG3*, *ERG6*, and *ERG11* orthologs from pathogenic fungi can complement the activity of their *S. cerevisiae* counterpart using a heterologous expression system

*ERG3, ERG6* and *ERG11* orthologs were identified from *S. cerevisiae* (*ScERG3*/*6*/*11*), *C. albicans* (*CalERG3*/*6*/*11*), *C. auris* (*CauERG3*/*6*/*11*), *N. glabratus* (*NgERG3*/*6*/*11*), *C. parapsilosis* (*CpERG3*/*6*/*11*) and *C. tropicalis* (*CtERG3*/*6*/*11*), along with additional *ERG11* orthologs *CYP51A* and *CYP51B* from *A. fumigatus* (*AfCYP51A*/*B*) using the Candida Genome Database (Lew-Smith et al. 2025), the Uniprot database (UniProt Consortium 2025) and FungiDB (Basenko et al. 2024) (Figure 1a). The *A. fumigatus* orthologs of *ERG3* and *ERG6* were omitted, as to our knowledge, no antifungal resistance mutation has been identified in these genes, outside of the Saccharomycotina clade. We calculated pairwise identities between the different orthologs (Figure 1b) using protein sequence alignments (Figure S1). As expected, more closely related species’ orthologs cluster together.

**Figure 1.**
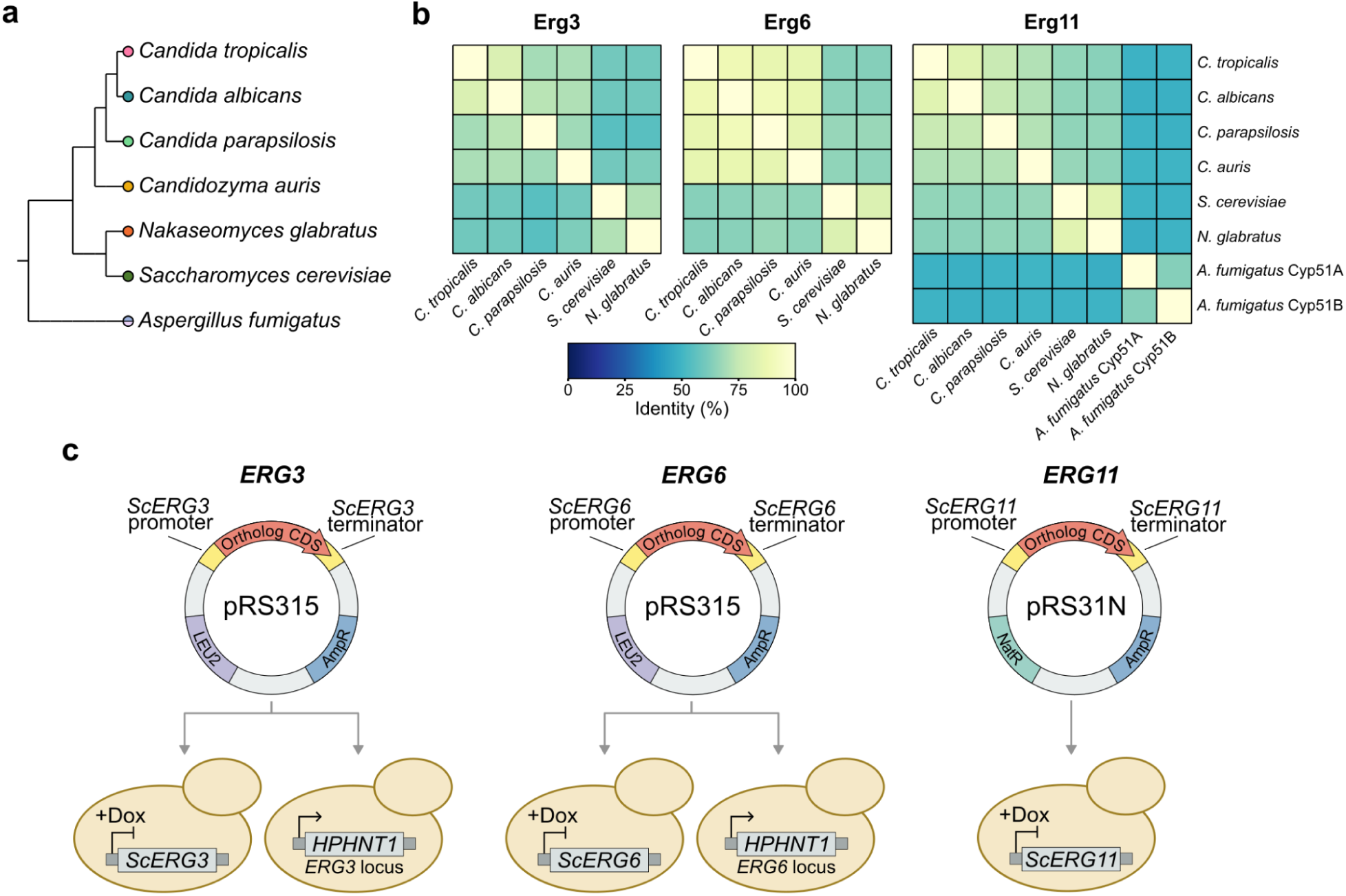
Plasmid-based heterologous expression system of orthologous genes *ERG3*, *ERG6* and *ERG11* in *Saccharomyces cerevisiae*. a) Cladogram of the fungal species from which orthologous sequences used in this study were obtained. Generated with PhyloT v2 based on the NCBI Taxonomy Database and visualized using iTOL (Letunic and Bork 2024). b) Comparison of protein sequence identity (%) between orthologs of Erg3, Erg6 and Erg11 calculated from the protein sequence alignment found in Figure S1. c) Codon-optimized *ERG3* and *ERG6* sequences were cloned into the pRS315 plasmid, then transformed into deletion (*erg3Δ*::*HPHNT1* and *erg6Δ*::*HPHNT1*) and repressible-expression (TetOpr-*ERG3* and TetOpr-*ERG6*) strains. Codon-optimized *ERG11* sequences were cloned into the pRS31N plasmid, then transformed into the repressible-expression strain TetOpr-*ERG11*.

The orthologous genes were expressed in *S. cerevisiae*. To do so in a systematic manner, a series of expression plasmids was constructed, all based on the pRS plasmid backbone. pRS-based plasmids are centromeric yeast plasmids, and should therefore maintain a single or low copy number (Sikorski and Hieter 1989). To keep wild-type like regulation, the promoter and terminator sequences of *S. cerevisiae* for each corresponding gene were used. To ensure proper gene translation, codon-optimized versions of the orthologous genes were synthesized. These were inserted into the plasmids, in frame with the corresponding promoter and terminator (e.g. *ERG3* orthologs with the *S. cerevisiae ERG3* promoter and terminator, and so forth) (Figure 1c, Left). The native *S. cerevisiae* genes were included as controls, using codon-optimized versions for *ERG3* and *ERG6*. Six *ERG3* plasmids, six *ERG6* plasmids, and eight *ERG11* plasmids were thus constructed. In total, a series of 20 plasmids were produced, allowing expression in *S. cerevisiae* of sterol synthesis genes from many of the most medically relevant pathogens. The proper construction of each plasmid was confirmed by whole plasmid sequencing.

*S. cerevisiae* strains were then constructed where the chromosomal copy of *ERG3*, *ERG6*, or *ERG11* was either deleted or had its expression repressed, leaving the plasmid gene copy as the only one being expressed. In doing so, both loss-of-function and gain-of-function mutational effects can be observed. For example, nonsense mutations in *N. glabratus ERG3* lead to resistance to multiple antifungal drugs (Lim et al. 2023), but expressing a nonsense mutant of *NgERG3* in a strain containing a wild-type allele of *ERG3* would not lead to an observable resistance phenotype. As *ERG3* and *ERG6* are not essential in *S. cerevisiae* (Gaber et al. 1989; Arthington et al. 1991), two deletion strains were generated using the *HPHNT1* selection marker (*erg3Δ*::*HPHNT1* and *erg6Δ*::*HPHNT1*). In contrast, *ERG11* is essential in *S*. *cerevisiae* (Kalb et al. 1987), so a simple deletion approach was not possible. Instead, a previously constructed strain was used (TetOpr-*ERG11*), using previously developed tools (Mnaimneh et al. 2004), where the chromosomal promoter was replaced by a Tet-repressible promoter, resulting in repressed expression of the chromosomal allele of *ERG11* in the presence of doxycycline down to levels that inhibit growth entirely (Bédard et al. 2024). The same strategy was used to generate ERG3 and ERG6 repressible strains. Our collection is thus composed of one deletion and one repressible-expression strain in which to measure the antifungal resistance of *ERG3* orthologs, one deletion and one repressible-expression strain for *ERG6*, and one repressible-expression strain for *ERG11* (Figure 1c, Right).

We first tested whether the orthologs were able to complement the function of the *S. cerevisiae* genes. As *ERG11* is essential in *S. cerevisiae*, complementation by the *ERG11* orthologs can simply be measured as growth. Since we found *ERG3* and *ERG6* to be essential in the presence of the surfactant SDS, complementation was tested by measuring growth in SDS containing media. Growth was evaluated in liquid media by measuring the optical density at 600 nm (OD) of cultures in 96-well plates over 48 hours. An empty plasmid (the pRS template plasmid) was used as a control for the absence of the ortholog, and the plasmids expressing *ERG3*, *ERG6* and *ERG11* sequences from *S. cerevisiae* were used as positive controls of complementation. For the *ERG3* and *ERG6* strains, 0.005% SDS was added to the growth medium, and for TetOpr strains, 10 µg/mL of doxycycline was added to repress expression of the chromosomal *ERG* gene copy.

All orthologs complement the loss of function of the *S. cerevisiae* native gene, as in all cases growth was similar to the *S. cerevisiae* positive control (Figure 2, Figure S2). In all strains, as expected, the empty plasmid showed no, or markedly low, growth. *NgERG3, CtERG6* and most notably *AfCYP51A* show a small but statistically significant (Tukey’s post-hoc HSD p < 0.05) decrease in growth compared to their respective *S. cerevisiae* gene. However, the growth of the three affected orthologs is more similar to their respective wild-type counterparts than to the control non-complementing plasmids, which show a marked difference with all orthologs.

**Figure 2.**
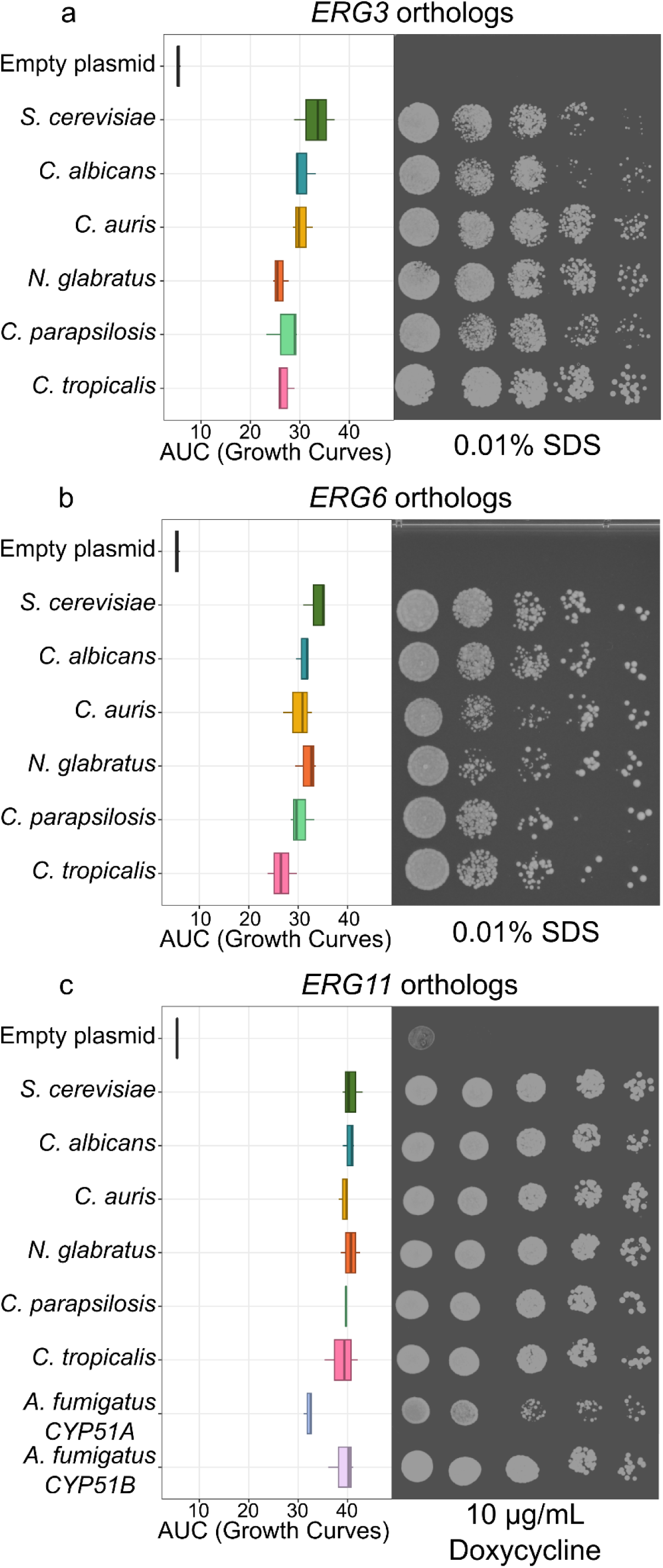
Complementation of orthologous *ERG* genes in *S. cerevisiae*. Complementation of the *ERG3* (a) and *ERG6* (b) *S. cerevisiae* genes with orthologs in the deletion backgrounds (*erg3Δ*::*HPHNT1* and *erg6Δ::HPHNT1*). Area under the curve (AUC) was calculated from growth curves in synthetic medium containing 0.005% SDS, and cultures spotted on solid synthetic medium containing 0.01% SDS. c) Complementation of the *ERG11* orthologs in the *S. cerevisiae* TetOpr-*ERG11* background. AUCs calculated from growth curves in synthetic medium containing 10 µg/mL of doxycycline to repress the chromosomal copy of *ERG11*, and cultures spotted on solid synthetic medium with 10 µg/mL doxycycline as well. Raw growth curves from which the AUCs were calculated are found in Figure S2. For all orthologs in all backgrounds, AUCs are calculated from 3 replicates.

In parallel, complementation was also tested using spot dilution assays on solid media (Figure 2). Once more, 10 µg/mL of doxycycline was used with the TetOpr-*ERG11* strain, while 0.01% SDS was added to solid media for the *ERG3* and *ERG6* complementation assays. As with the growth curves, all orthologs grow similarly to the *S. cerevisiae* positive control, demonstrating full complementation. As the growth of all orthologs is easily differentiable from the empty plasmid (negative control), we can use the heterologous expression system to measure both gain-of-function and loss-of-function mutations.

For the TetOpr-*ERG3* and TetOpr-*ERG6* strains, we did not achieve full inhibition of the chromosomal copy of the *ERG* gene in liquid media with the same concentration of 0.005% SDS (Figure S3). We tested concentrations up to 0.0125% SDS, where there was marked difference between the growth of the negative control and the complementing orthologs (Figure S3). However, the difference is clearer in the deletion strains, and these will be used going forward, though the TetOpr strains can still be useful when conditional repression is necessary.

### Phenotypic effects of resistance-conferring mutations are recreated in the heterologous expression system

Next, we tested whether the heterologous expression system could be used to recreate the phenotypes of known resistance-conferring mutations. The FungAMR database (Bédard et al. 2025) was queried for examples of single amino acid changes in orthologs of *ERG3*, *ERG6*, and *ERG11*, which would confer resistance to an antifungal drug. For *ERG3*, G111R was shown to confer resistance to micafungin in *C. parapsilosis* (Rybak et al. 2017; Hartuis et al. 2024). For *ERG6,* G127R was identified as a result of experimental evolution for nystatin resistance in *S. cerevisiae* (Gerstein et al. 2012). *ERG11* has numerous reports of resistance mutations, and the K143R mutation in *CalERG11* was selected as it was the first identified in an azole-resistant clinical isolate (Manavathu et al. 1999; Perea et al. 2001). In FungAMR, this same mutation is also reported to increase resistance to azoles in *CauERG11*, *CpERG11* and *CtERG11* (Bédard et al. 2025). As *AfCYP51A* and *AfCYP51B* have not been found to be resistant through K143R (or corresponding mutations), a mutation known to confer resistance in *A. fumigatus* was added. We selected G448S in *AfCYP51A,* a mutation first observed in a clinical isolate which shows resistance to many azoles (Howard et al. 2009). The sequence alignment (Figure S1) reveals that the corresponding mutation in *AfCYP51B* is G457S, which has also been shown to be resistant to voriconazole (Handelman and Osherov 2023), meaning that putative resistance in both paralogs from this mold can be tested this way.

To recreate the aforementioned mutations of known effect, we used a standard site-directed mutagenesis approach based on reverse-complement primers encoding the mutation of interest, that can easily be applied to any mutation. We used spot dilution assays on media containing various antifungal drugs to assess antifungal resistance in the wild-type and constructed mutant orthologs. In all cases, the empty plasmid background not expressing an *ERG* gene, and the wild-type orthologous sequence were included as controls to compare against the mutant phenotype.

We applied spots of wild-type *CpERG3* and *CpERG3* G111R on media containing 0.025 µg/mL of micafungin (Figure 3a). More growth is observed for the *CpERG3* G111R mutant compared to wild-type, demonstrating that the heterologous expression system can recreate known resistance phenotypes. Interestingly, the *ERG3* deletion strain containing the empty plasmid also grows in the presence of micafungin. Deletion of *ERG3* has been shown to confer resistance to caspofungin, another echinocandin class drug (Markovich et al. 2004; Bhakt et al. 2022). Furthermore, *ERG3* nonsense mutations in *N. glabratus* and *C. albicans* have led to resistance to various echinocandins (Spettel et al. 2019; Lim et al. 2023).

**Figure 3.**
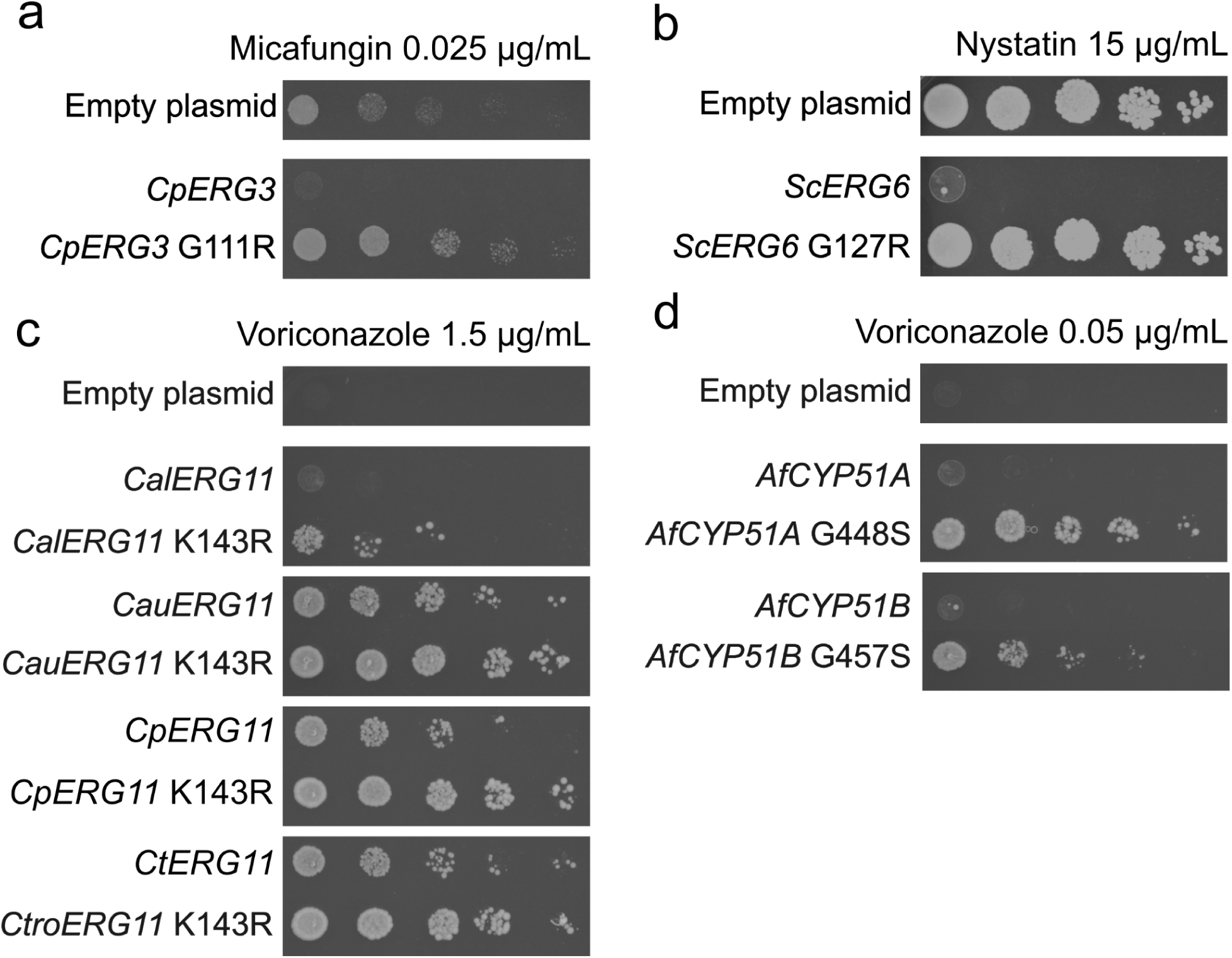
Resistance-conferring mutations from different orthologs also confer resistance when they are recreated and expressed in *S. cerevisiae*. Spot dilution assays showing susceptibility, or lack thereof, of empty plasmids, wild-type and mutant orthologs of a) *CpERG3* to 1.5 µg/mL of voriconazole, b) *ScERG6* to 15 µg/mL of nystatin, c) multiple orthologs of *ERG11* to 3 µg/mL of voriconazole and d) *AfCYP51A/B* to 0.05 µg/mL of voriconazole.

The resistant phenotype of *ScERG6* G127R was also measured using a spot dilution assay, this time on 15 µg/mL of nystatin (Figure 3b). As expected, the cells expressing the wild-type gene did not grow while those expressing the mutant gene did. Once more, the cells containing the empty plasmid grew, which confirms previous observations that *ERG6* deletion and nonsense mutations can lead to nystatin and amphotericin B resistance in *S. cerevisiae* (Gerstein et al. 2012; Bhattacharya et al. 2018), a phenotype that was also observed in *C. auris* (Carolus, Sofras, Boccarella, Sephton-Clark, et al. 2024; Carolus, Sofras, Boccarella, Jacobs, et al. 2024).

All 4 tested mutants of *ERG11* yeast orthologs and the 2 *A. fumigatus* orthologs also recreated the known resistance phenotype. Spot dilution assays for yeast orthologs were done on 1.5 µg/mL of voriconazole (Figure 3c), while the *A. fumigatus* orthologs were spotted on 0.05 µg/mL (Figure 3d). As previously, solid media supplemented with 10 µg/mL of doxycycline for the TetOpr-*ERG11* strain was used to repress expression of the chromosomal copy of *ERG11*, making growth dependent on the gene expressed from the plasmid. In this case, the cells containing the empty plasmid did not grow and are non-viable, as expected since *ERG11* is an essential gene and it was not expressed from either the chromosome or the plasmid.

In short, all the tested mutations confer resistance to various antifungal drugs in our heterologous expression system, as observed in other studies, while the expression of the corresponding wild-type gene does not allow growth. This validates the heterologous expression system as a relatively simple and rapid method for testing the resistance-conferring potential of mutations.

### The heterologous expression system allows resistance testing of novel mutations

As the heterologous expression system demonstrated its ability to recreate known resistance phenotypes, we next used it to measure the phenotypes of mutations of unknown effect. As a starting point, we recreated the *ERG3* G111R, *ERG6* G127R and *ERG11* K143R mutations in the orthologs for which the effect of these mutations had not been previously characterized. Corresponding positions across orthologs were identified (Figure S1), and the equivalent mutations were created across all orthologs of the same gene. Once more, resistance to micafungin, nystatin or voriconazole was measured using spot dilution assays.

Surprisingly, mutations equivalent to the G111R mutation in other *ERG3* orthologs do not show any resistance to micafungin as the *C. parapsilosis* mutant does (Figure S4a). In fact, only cells containing the empty plasmid or expressing *CpERG3* G111R grow in spot dilution assays with 0.025 µg/mL of micafungin. As *ERG3* mutations have been previously linked to azole resistance (Bédard et al. 2025), we also tested all wild-type and mutant orthologs in spot dilution assays on 1.5 µg/mL of voriconazole (Figure 4a). Here, all mutants show a resistant phenotype, along with wild-type *CpERG3* (Figure S4b). *C. parapsilosis* is not known to be inherently azole resistant (Escribano and Guinea 2022). Further investigation, using the heterologous expression system or in *C. parapsilosis* will be necessary to elucidate this differential phenotype.

**Figure 4.**
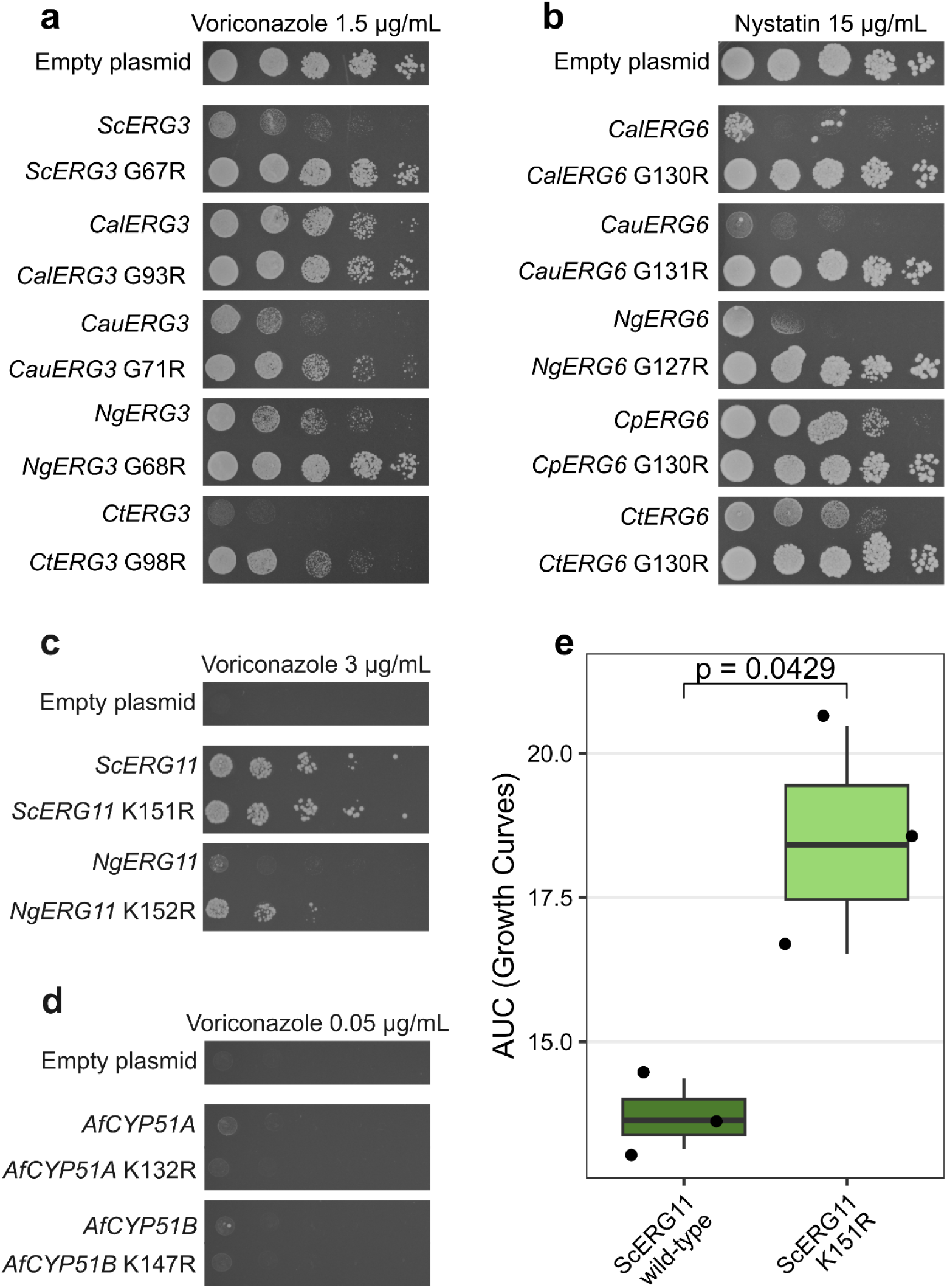
Spot dilution assays and growth curves with the heterologous expression system can measure the resistance of mutations of unknown effect Spot dilution assays showing susceptibility, or lack thereof, of empty plasmids, wild-type and mutant orthologs of a) multiple orthologs of *ERG3* to 1.5 µg/mL of voriconazole, b) multiple orthologs of *ERG6* to 15 µg/mL of nystatin, c) Sc*ERG11* and *NgERG11* to 3 µg/mL of voriconazole and d) *AfCYP51A/B* to 0.05 µg/mL of voriconazole. e) Area under the curve (AUC) measurements of growth of *ScERG11* WT and the K151R mutant, in the presence of 2 µg/mL of voriconazole. Two-tailed Welch’s t-test shows a significant difference between the AUC of the variants (p=0.0429).

On the other hand, while all wild-type alleles of *ERG6* are sensitive to 15 µg/mL of nystatin, all tested mutant alleles confer higher growth on the same concentration of nystatin (Figure 4b). The resistance-conferring potential of the G127R mutation is thus conserved across all tested species.

Additionally, *ERG11* orthologs for which resistance phenotype had not be assessed, *ScERG11* and *NgERG11,* along with both *A. fumigatus* orthologs, were tested for growth on voriconazole, and show opposite effects (Figure 4c, d). The two orthologs most closely related to the *Candida* orthologs, *ScERG11* and *NgERG11*, both show resistance phenotypes. For *ScERG11* the effect is extremely subtle, as the wild-type allows growth on solid media, even in the presence of 3 µg/mL of voriconazole. However, growth in liquid media containing 2 µg/mL of voriconazole shows a statistically significant difference (two-tailed Welch’s t-test, p=0.0429) between the AUC of growth of wild-type *ScERG11* and *ScERG11* K151R (Figure 4e). The small resistance advantage of the mutant is caused more by the low intrinsic susceptibility of wild-type ScERG11, as shown by the high growth in the spot dilution assay, than by a lack of resistance of the mutant. Nevertheless, *NgERG11* presents a clearer pattern, as the mutant grows in spots on 3 µg/mL of voriconazole, while the wild-type does not. However, in the case of the two *A. fumigatus* orthologs, both the wild-type and mutant alleles show no growth, even in the presence of a low (0.05 µg/mL) concentration of voriconazole. *A. fumigatus* orthologs are less closely related to (Figure 1a) and share less sequence identity with (Figure 1b) the other *ERG11/CYP51* orthologs. Furthermore, no resistance-conferring mutations have been reported at the mutated positions in Af*CYP51A* or *AfCYP51B* in the FungAMR database (Bédard et al. 2025). Thus, a differential effect of a mutation between yeast and mold orthologs has been demonstrated using the heterologous expression system.

### Use of the heterologous expression system for the discovery of novel resistance-conferring mutations

Beyond testing specific reported point mutations, the plasmid collection could also be used to create and screen large numbers of mutations at once. Previously, a heterologously expressed *C. albicans ERG11* deep mutational scanning library was used to assay all possible mutations near the active site of Erg11p (Bédard et al. 2024). Here, the heterologous expression system capacities are further demonstrated by discovering new resistance-conferring mutations using random mutagenesis.

We targeted the region of *CalERG11* from codons 373 to 520, roughly corresponding to fragment 4 in (Bédard et al. 2024). This region covers 63 positions that had been systematically mutated and characterized, while 85 positions had not been phenotyped. Targeting this fragment thus allowed us to randomly recreate mutations of known effect, which act as controls, as well as novel mutations that have never been assessed. To construct the variant library, we amplified the fragment using an error-prone polymerase, then cloned it into the expression plasmid, and transformed into the TetOpr-ERG11 strain (Figure 5a). This created a heterologous expression library containing single or multiple different *CalERG11* mutations. The library was then grown in liquid cultures containing 0.08 µg/mL of voriconazole for approximately four generations, and subsequently plated on solid media containing 0.4 µg/mL of voriconazole, thus enriching resistant mutants (Figure 5b). A preliminary assay using resistant and wild-type control strains showed a ∼3.5-fold enrichment of the resistant strain (see methods). The colonies enriched for resistance were pooled and sequenced to identify resistance mutations based on their enrichment in the final sample.

**Figure 5.**
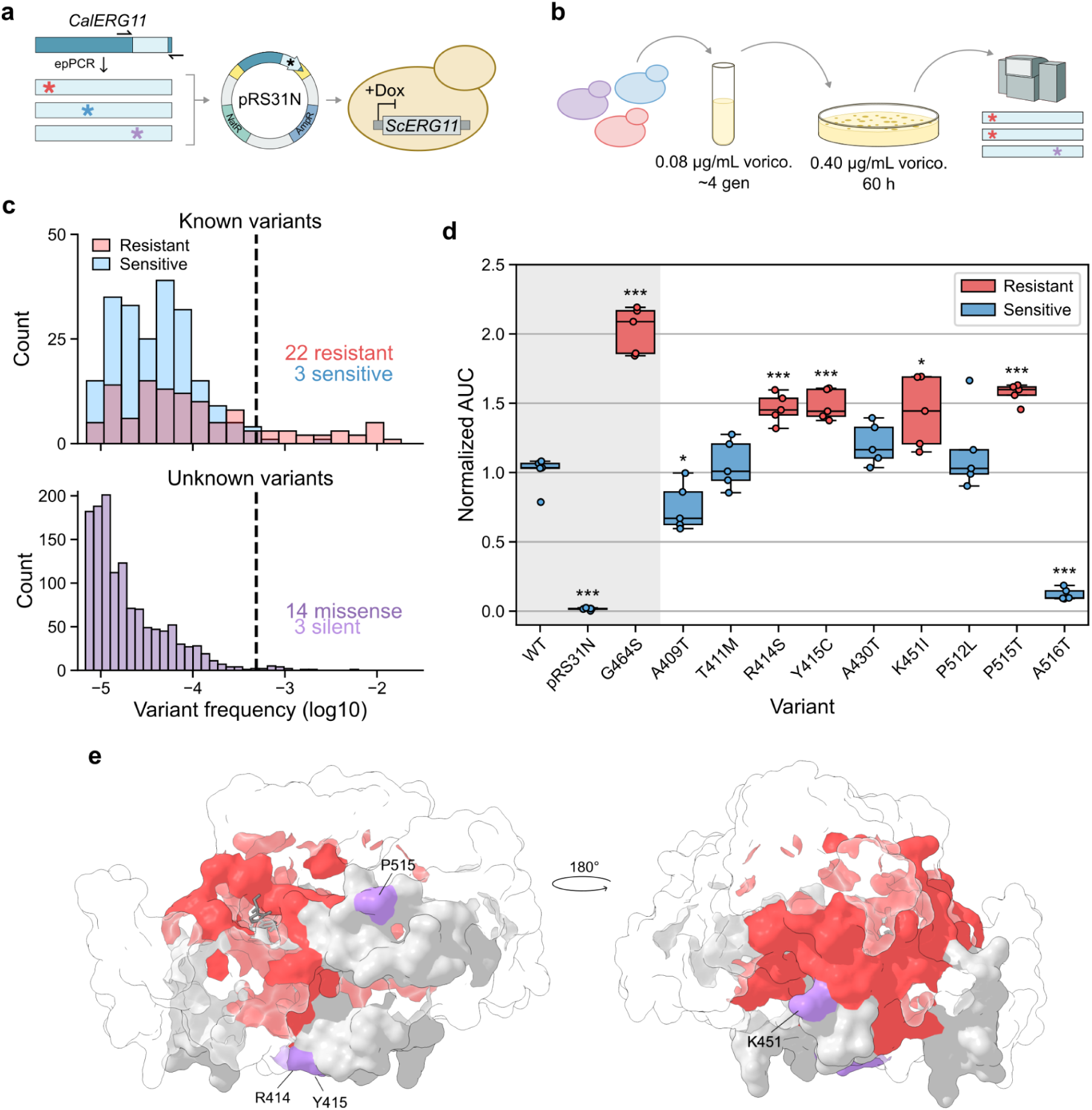
Discovery of four novel voriconazole resistance-conferring mutations in *CalERG11*. a) A region of *CalERG11* spanning nucleotide positions 1118-1559 (codons 373-520) was randomly mutated by error-prone PCR. The mutated *CalERG11* fragments were cloned in the pRS31N plasmid containing the rest of the *CalERG11* sequence and expressed in the TetOpr-*ERG11* strain. b) The variant library was grown in a liquid culture containing 0.08 µg/mL voriconazole (labelled vorico. on figure) for 4 generations and plated on solid media containing 0.4 µg/mL voriconazole to select for resistant variants. c) Distribution of variant frequency after selection for the variants previously characterized in Bédard et al., 2024 (top) and the variants that were not previously characterized (bottom). Variant frequency is the median of 2 to 4 selection experiments and of combined synonymous codons when available. Black dotted line represents the 99th percentile of the frequency of sensitive variants, which was used as a threshold value for the selection of variants to validate. d) Growth in 0.086 µg/mL voriconazole of variants selected for validation (n = 5 replicates). Areas under the curve (AUC) calculated from growth curves were normalized relative to the wild-type (WT). Experiment controls are in the gray shaded area and include the known highly resistant variant G464S (Bédard et al. 2024) and the empty plasmid control (pRS31N). Four previously uncharacterized variants grew more than the wild-type and were classified as resistant (two-sided independent t-test with correction for multiple comparisons using the Benjamini-Hochberg method; * : P < 0.05, *** : P < 0.001). Variants with similar or lower growth than the wild-type were classified as sensitive. e) Voriconazole resistance mutations mapped on the CalErg11p protein structure (PDB ID : 5V5Z). Positions where at least one mutation conferred resistance in Bédard et al., 2024 are in red. Positions of new mutations identified in this study are in purple. Positions included in the random mutagenesis library for which no resistance mutations were identified are in grey.

Of the 1,615 variants retrieved after selection, 326 had previously been characterized as either sensitive or resistant to voriconazole (Bédard et al. 2024). We found a correlation (two-tailed Spearman’s rank correlation, ρ = 0.62, p = 1.31 x 10^-115^) between the frequency of mutants in the initial library and after selection, suggesting that final mutant frequencies largely remain dependent on their initial proportions even if selection has enriched for resistant mutants (Figure S5). Known resistant mutations clearly departed from the correlation, being much more frequent after selection, along with a number of unknown mutants (Figure S5). To more systematically identify enriched resistant mutants, we set a threshold at the 99th percentile of the frequency of known sensitive mutants to classify our newly discovered mutants as resistant or sensitive. This threshold is not perfect, as it excludes known resistant mutations and includes a few known sensitive mutations, but it sets a limit beyond which resistant mutations should be much more frequently found. Encouragingly, the strongest resistance phenotypes measured in this screen are for mutations that were previously known to cause high-resistance levels (for instance G464S) (Figure S5), suggesting that most strong-effect mutations among these positions may already be known but would have been discovered using this approach.

Past the assigned threshold, we identified 17 mutations outside the previously characterized positions (Bédard et al. 2024), 14 of which being missense mutations (Figure 5c). We chose 9 mutants that had only a single codon substituted, and rebuilt them using the same site-directed method as described above. Their growth was measured in liquid media containing 0.086 µg/mL of voriconazole, along with 3 controls (Figure 5d). A two-tailed t-test (p < 0.05) identified 4 as being resistant compared to the wild-type. This result suggests that a fraction of mutants growing on the selection plates are not resistant, but are present after selection because of their high frequency in the initial pool of mutants (Figure S5). Mapping the position of the newly discovered resistance-conferring mutations to the structure of *CalERG11* shows that their positions are contiguous to previously known sites of resistance-conferring mutations and thus relatively close to the active site (within 22Å of the heme) (Figure 5e). Their positioning suggests that the mode of action of the newly discovered mutations is similar to other mutations near the active site (Lupetti et al. 2002; Bédard et al. 2024). In short, the heterologous expression system can also be used in conjunction with simple methods, such as random mutagenesis, to screen for novel resistance-conferring mutations.

## Discussion

Antifungal resistance is a worldwide challenge, and only continues to grow. To better tackle this problem, a more thorough understanding of the effects of various mutations in key genes is necessary. Point mutations, such as in genes coding for enzymes involved in the sterol synthesis pathway, are often found in resistant isolates (Robbins et al. 2017; Revie et al. 2018; Lee et al. 2023). Validating the involvement of these mutations in resistance, rather than their fortuitous co-occurrence, is key for the interpretation of sequencing data. The precise consequences of such mutations can be slow or difficult to study due to the limitations of the current genetic tools available for fungal pathogens. Here, we show the potential of standardized systems for the identification of resistance-conferring mutations.

Our approach is based on a large body of previous work. Using heterologous expression to study genes from fungal pathogens has a long history, and has led to many scientific advances, including confirming the causal link between mutations observed in clinical isolates and their resistance phenotype (Prasad et al. 1995; Cools et al. 2010; Xiang et al. 2013). Validating the causal role of mutations is critical in the field, but it is rarely done, as shown by the numerous entries of resistance mutations in databases that have no experimental validation (Bédard et al. 2025). We extended previous work and constructed a series of plasmids containing 6 *ERG3* and *ERG6* orthologs as well as 8 *ERG11* orthologs from clinically relevant fungal pathogens, all of which complementing the function of the *S. cerevisiae* genes. The expected phenotype was observed when introducing known resistance-conferring mutations, confirming the usefulness of this approach for testing the phenotypic effects of mutations.

In addition to validating the impact of mutations found in fungal pathogens, our plasmids can be combined with high-throughput methods of library construction to screen large numbers of substitutions and discover mutations not yet reported in pathogens. For example, deep mutational scanning of *CalERG11* led to the experimental validation and discovery of hundreds of azole resistance mutations (Bédard et al. 2024). A potential alternative to deep mutational scanning for discovering resistance mutations is random mutagenesis. Here, we used random mutagenesis to construct a mutant library of a segment of *CalERG11* that was subjected to deep mutational scanning, but that also included 85 positions not previously examined. Our screen identified 4 mutations conferring partial resistance to voriconazole (R414S, Y415C, K451R, and P515T), which, to our knowledge, have not previously been associated with azole resistance in *C. albicans* or other fungal pathogens. Although P515S was previously observed with two other mutations in a fluconazole-resistant *Candida dubliniensis* isolate, the causal effect of this mutation alone on resistance was not assessed (Perea et al. 2002). Overall, these 4 identified and validated mutations are located relatively close to the active site, and close to previously known resistance-conferring mutations, suggesting a similar mechanism of resistance.

While random mutagenesis is less technically demanding and expensive than deep mutational scanning, it comes with limitations. For instance, in our experiments, variant frequencies were highly skewed, with some variants being more than 100x more frequent than others. This bias affected our ability to separate resistant from sensitive variants solely based on their frequency after selection, which does not eliminate all susceptible WT and mutants at the drug concentrations used. Nevertheless, for some applications in which deep mutational scanning is not affordable, for instance, the comparison of multiple orthologs in parallel, random mutagenesis, and expression in *S. cerevisiae* remain of interest. The protocol could be improved by increasing the ratio of true positives at the end of screening, for instance by augmenting drug concentrations or the number of selection generations. However, one must balance false positives and false negatives, as stronger selection may prevent the discovery of resistance mutations with weaker effects.

The plasmids we constructed will be made available through the Addgene repository, making the system accessible to the scientific community. The system is also easily expandable, as new plasmids containing orthologs from other species can be constructed using Gibson assembly, with any of the existing plasmids as a template. In short, the approach is now standardized, expanded, and available to the scientific community for easy use.

## Methods

### Growth conditions

Bacterial cultures were grown in liquid 2xYT broth medium (See Table S1 for details of culture media used). When making solid media, 2% agar was added. When necessary, 100 µg/mL of ampicillin (Amp, Bioshop Canada), was added as well. Yeast cultures were grown in synthetic complete (SC) medium, with the pH buffered to 6.0. When working with cells containing pRS315-derived plasmids (plasmids expressing *ERG3* and *ERG6* orthologs) the medium was made using -leucine drop-out amino-acid mix, to allow for selection of the auxotrophic *LEU2* marker on pRS315 (Sikorski and Hieter 1989). For selection of the pRS31N-derived plasmids (Hénault et al. 2024) (plasmids expressing *ERG11* orthologs), 100 μg/mL of nourseothricin (Nat, Jena Bioscience) was added. For selection of the *erg3Δ*::*HPHNT1* and *erg6Δ*::*HPHNT1* strains, 250 µg/mL of hygromycin B (HygB, Bioshop Canada) was added. When specified, 10 µg/mL of doxycycline (Dox, Bioshop Canada) or the specified concentration of sodium dodecyl sulfate (SDS, Bioshop Canada) were added. Also when specified, voriconazole (TCI America), nystatin (BioShop Canada), or micafungin (Toronto Research Chemicals) were diluted in DMSO (BioShop Canada) and added in the specified quantities. Yeast growth was done at 30°C with the addition of shaking at 250 rpm for growth in liquid medium.

### Gene sequences

The gene sequences of *ERG3, ERG6,* and *ERG11* for different species were obtained from various databases. The *S. cerevisiae* gene sequences were obtained from the S288C strain in the Saccharomyces Genome Database (Engel et al. 2025). Gene sequences for *C. albicans, N. glabratus, C. parapsilosis* and *C. auris* were obtained from the respective reference genomes from the Candida Genome Database (Lew-Smith et al. 2025). *C. tropicalis* and *A. fumigatus* gene sequences were obtained from Uniprot (UniProt Consortium 2025) and cross-referenced using FungiDB (Basenko et al. 2024). All details of gene and protein sequences are available in Table S2. For *ERG3* and *ERG6,* gene sequences from CTG clade fungi (*C. albicans, C. auris, C. tropicalis and C. parapsilosis*) were inspected, and CTG codons were changed to TCT codons, which code for serine in *S. cerevisiae*. All sequences were then codon optimized for expression in *S. cerevisiae* using the GenSmart Codon Optimization Tool, with *ERG11* orthologs optimized directly from protein sequences (GenScript Biotech Corporation).

### Sequence alignments

Protein sequences were aligned using the MAFFT webserver, with the G-INS-i method (Katoh and Standley 2013; Katoh et al. 2019). A fasta file of all protein sequences of either Erg3p, Erg6p or Erg11p was used as input. The options correspond to the command “mafft --inputorder --leavegappyregion --maxiterate 2 --retree 1 --globalpair <input>”. From these alignments, percent identity scores were calculated using the “seqidentity” command from the R package bio3d (Grant et al. 2006) (https://github.com/Landrylab/Jordan-et-al-2026). Sequence alignments were visualized using Jalview 2.11 (Waterhouse et al. 2009).

### Cloning

The codon optimized sequences were synthesized (GenScript). We added 20 nucleotides of homology at the 5’ and 3’ ends, which match to the promoter and terminator of the corresponding gene in *S. cerevisiae* for *ERG3* and *ERG6*. Plasmid construction was based on previous work (Bédard et al. 2024). pRS31N-CalbERG11 and pRS31N-ScerERG11 had already been built for this previous project. Template plasmids were constructed containing only the promoter and terminator of *S. cerevisiae ERG3*, *S. cerevisiae ERG6,* or *S. cerevisiae ERG11*. Procedures varied slightly between *ERG3* and *ERG6* orthologs compared to *ERG11* orthologs. The promoter and terminator were respectively defined as the 1000 bp preceding and the 200 bp following the gene for *ERG3* and *ERG6*. For *ERG11* the native promoter was 878 bp preceding the gene and the terminator was 319 bp following the gene (Bédard et al. 2024). The *ERG3* and *ERG6* plasmids were based on pRS315, which contains the *LEU2* auxotrophic marker (Sikorski and Hieter 1989). For *ERG11*, the plasmid used is pRS31N (Hénault et al. 2024) which contains the NATNT2 resistance marker. All plasmids were amplified outwards from the insertion site (the junction of the promoter and terminator), using a series of primers that added homology to the different *ERG3, ERG6, and ERG11* orthologs. See Table S3 for all primers used. The amplified plasmid backbones were then assembled with the codon optimized ortholog sequences using Gibson assembly (New England Biolabs, Inc. 2020), and transformed into *E. coli*. The cloning resulted in orthologous genes that are in frame with the *S. cerevisiae* promoter and terminator. Proper plasmid construction was verified using whole plasmid sequencing (Plasmidsaurus and Flow Genomics). Plasmids were extracted from *E. coli* using the Presto Mini Plasmid Kit (Geneaid).

The construction of the mutant orthologs was started from the wild-type gene plasmids, and was done using site-directed mutagenesis, based on the QuickChange Site-Directed Mutagenesis System (Stratagene), and previously published work (Bachman 2013). PCR products were digested with 20 U of DpnI (New England Bioscience) at 37 °C for 90 minutes to remove the template plasmid (which does not contain the mutation). These digested products were then transformed into *E. coli* strain MC1061, for ligation and selection. Positive transformants were selected using ampicillin, as the pRS plasmids all contain an ampicillin selection marker. The presence of mutations was verified using Sanger sequencing.

When testing the *ERG11* orthologs, we realized that the reference sequence used for *CpERG11* contains a phenylalanine at position 132, instead of a tyrosine, which is a substitution known to confer azole resistance (Hartuis et al. 2024). Many known wild-type alleles of *CpERG11* have a tyrosine at position 132 (Berkow et al. 2015; Arastehfar et al. 2020), including the ATCC type strain 22019 (Benton et al. 2021). Using the same aforementioned site-directed mutagenesis method, the phenylalanine was mutated to a tyrosine before the construction of other mutations, thus allowing measurement of resistance phenotypes. Similarly, a mutation in codon 227 leading to a Y227H amino acid substitution was identified in *ScERG11* and was reverted to the wild-type nucleotide by site-directed mutagenesis.

### Yeast strain construction

The *S. cerevisiae ERG11* doxycycline-repressible promoter strain (TetOpr-*ERG11*) was previously constructed and used in another work, where it was named *ScERG11*-DOX, (Bédard et al. 2024), and followed the scheme of the Yeast Tet-Promoters Hughes collection (Mnaimneh et al. 2004). TetOpr-*ERG3* and TetOpr-*ERG6* were constructed in the same way as TetOpr-*ERG11*, replacing the endogenous promoter with the doxycyclin-repressible TetO_7_ promoter (referred to as TetOpr throughout the text). For the *ERG3* and *ERG6* deletion strains (*erg3Δ*::*HPHNT1* and *erg6Δ*::*HPHNT1*), the HPHNT1 cassette was amplified from the pFA6-HPHNT1 plasmid (Janke et al. 2004) with primers adding homology to the promoter and terminator of either *ERG3* or *ERG6*. These deletion cassettes were transformed into BY4741 yeast strain (*MATa his3Δ leu2Δ met15Δ ura3Δ*), using a standard lithium acetate yeast transformation (Gietz and Schiestl 2007). Transformants were selected on media containing HygB, and the transformation of the deletion cassette was verified using Sanger sequencing.

Transformation of control, wild-type and mutant plasmids into the TetOpr and deletion strains was done slightly differently, depending on the gene expressed. For the *ERG3* and *ERG6* plasmids, a DTT based transformation protocol was used which was modified from (Elble 1992), as this was found to be more efficient. Briefly, precultures were grown overnight, then 1.5 mL of culture was centrifuged at 21 000 x g for 1 minute in a microcentrifuge tube. The supernatant was decanted, leaving about 100 µL of liquid along with the cell pellet. Then, 5 µL of boiled and subsequently cooled 10 mg/mL ssDNA along with 250-2500 ng of miniprepped plasmid were added, and then mixed. Subsequently, 0.5 mL of sterile PLATE mixture (18 % PEG 3350 (BioShop Canada), 100 mM lithium acetate (Thermo Scientific), 10 mM Tris-HCl (BioShop Canada), and 1 mM EDTA (BioShop Canada)) was added, and then mixed. Next, 20 µL of DTT 1 M (BioShop Canada) was added, and the tube was mixed once more. The mixture was allowed to incubate at room temperature for 6 hours, then incubated for 10 minutes at 42 °C. Finally, 100 µL of this mixture was plated on selective SC -leucine plates, and the plates were incubated at 30 °C for 72 hours. For the *ERG11* plasmids, a standard lithium acetate yeast transformation was used (Gietz and Schiestl 2007). Transformants were selected using Nat resistance.

### Growth curves

Growth in liquid media was measured to test the complementation of the different gene orthologs. First, overnight cultures were inoculated in SC media by picking a single colony from a Petri dish without leucine or containing Nat to select either the pRS315 or pRS31N based plasmids. For TetOpr-*ERG3* and TetOpr-*ERG6*, 10 µg/mL of Dox were added at this point. From these overnight cultures, growth curves were inoculated in 96 well plates. The overnight cultures were diluted to a final concentration of 0.01 optical density at 600 nm (OD) in 200 µL of SC media, maintaining leucine or Nat selection. For TetOpr-*ERG3* and TetOpr-*ERG6*, preliminary tests showed incomplete inhibition of gene expression, and so overnight precultures were diluted to 0.2 OD in SC media with 10 µg/mL of Dox. These cultures were allowed to grow for 6 hours, and were then diluted in the same manner as the precultures. For the next step, for the TetO promoter strains, 10 µg/mL of Dox was added. For the *ERG11* orthologs, nothing else was added, while for the *ERG3* and *ERG6* orthologs, either 0.005%, 0.0075%, 0.01% or 0.0125% of SDS was added. For the deletion strains, HygB was added, along with 0.005% SDS. Each strain was grown in three replicates in different wells.

Growth was measured in Agilent Biotek Epoch 2 plate readers (Agilent). 96-well plates were incubated at 30 °C for 48 hours without shaking, and the OD was measured every 15 minutes. The analysis of the growth curves was done using a custom R script. The tidyverse collection of packages was used for analysis (Wickham et al. 2019). Significant differences in the area under the curve (AUC) of the growth curves were calculated by fitting an ANOVA using R 4.5.1, and then executing Tukey’s HSD test with an alpha of 0.05 using the R package agricolae version 1.3-7 (de Mendiburu 2023).

Liquid growth was also measured to distinguish the susceptibility to voriconazole of wild-type *ScERG11* and *ScERG11* K151R. Growth was measured as described above for complementation. However, 2 µg/mL of voriconazole was added to the medium. AUC was calculated using the same custom script. The difference in AUC was evaluated using a two-tailed t-test with an alpha of 0.05, which was implemented using R 4.5.1.

### Spot dilution assays

Complementation and resistance were measured using spot dilution assays on solid media. Overnight cultures of cells were grown in SC media, with leucine selection or Nat selection for pRS315 or pRS31N containing cells, respectively. These precultures were diluted to 1 OD in SC media. Four subsequent dilutions of 1/5 were done in SC media. Droplets of 4 µL of each dilution were spotted on solid plates, except for plates containing SDS, where only 2 µL was spotted (due to lowered surface tension causing merging of spots). Different concentrations of SDS, Dox, HygB, voriconazole and nystatin were added, as detailed in the results section. Spotted plates were incubated at 30 °C for 72 hours, then photographed using one of two automated photography platforms (spImager or BM5-SC1, S&P Robotics Inc.).

### Random mutagenesis plasmid libraries construction

The random mutagenesis of the *CalERG11* region from nucleotide positions 1118 to 1559 (amino acid residues 373 to 520, roughly corresponding to fragment 4 (F4) in Bédard et al., 2024) was performed using the GeneMorph II Random Mutagenesis Kit (Agilent). The *CalERG11* region was amplified from a pRS31N-CalERG11 plasmid where the ampicillin resistance marker had been replaced with a kanamycin resistance marker. Briefly, the KANMX cassette was amplified from the pUG6 plasmid (Güldener et al. 1996) with primers adding homology to the AmpR promoter and plasmid backbone. The plasmid backbone was amplified outwards of the insertion site and the amplified KANMX cassette was then inserted using Gibson assembly.

The random mutagenesis PCR reaction was performed following the kit’s instruction for the preparation of the sample mix and the cycling parameters. PCR amplifications were done with 500, 650, 700 and 750 ng of target DNA (7.9, 10.3, 11.1 and 11.8 µg of total DNA) to achieve a low mutation frequency. The PCR products were purified using magnetic beads (Axygen AxyPrep Mag PCR Clean-up kit) in two steps. First, 24 µL of magnetic beads were added to 48 µL of the PCR product and the supernatant, containing only the smaller DNA fragments, was collected. The supernatants containing the PCR products were then digested with 20 U of DpnI (New England Biosystems) at 37 °C for 60 minutes. In the second step, 10 µL of magnetic beads were added to the digested supernatant, which was then washed twice with 100 µL of 80% ethanol before being dried and resuspended in 15 µL of Tris 10 mM pH 8.0. We were not able to get rid of all the parental DNA with those purification steps, as bands with a length corresponding to the template plasmid could still be seen on an agarose gel. However, the plasmids used as template for the random mutagenesis PCR and as the backbone for the subsequent Gibson assembly have different selection markers, reducing carry-on of template DNA in the library.

We constructed a variant pRS31N-CalERG11 plasmid, where a NdeI restriction site was added in *CalERG11* F4 by site-directed mutagenesis (mutation C1381G), using the site-directed mutagenesis method described above. This plasmid was used as the backbone for the cloning of the purified random mutagenesis PCR products. Before cloning, the pRS31N-CalERG11 vector (2 µg) was first digested with 40 U of NdeI (New England Biolabs) in 4 µL of CutSmart Buffer (10X) and water (total volume of 40 µL) at 37°C for 2 h and then inactivated at 65°C for 20 minutes. The digested backbone was then amplified by PCR with primers pointing outwards of *CalERG11* F4, digested with 20 U of DpnI (New England Biolabs) at 37 °C for 1 h and purified using magnetic beads (Axygen AxyPrep Mag PCR Clean-up kit). Cloning of the mutagenesis PCR products was done following Gibson assembly protocol with 50 ng of the plasmid backbone and 33 ng of the insert. Transformants were selected using ampicillin resistance. Following transformation, 5 mL of 2xYT medium was added to the plates and colonies were scraped using a sterile glass rake. At least 10,000 colonies were recovered for each cloning. Plasmids were extracted from a cell pellet corresponding to an OD of 25, using the Presto Mini Plasmid Kit (Geneaid). We thus constructed four plasmid libraries from independent random mutagenesis PCR reactions, referred to as libraries F4-500, F4-650, F4-700 and F4-750. Aliquots from the colony scrapings were stored at -80°C in 15% glycerol.

To validate the proper ligation of the mutated fragment in the plasmid backbone, two plasmids per library, extracted from isolated colonies from streaking the glycerol stocks, were sent to whole plasmid sequencing (Flow Genomics). One plasmid from library F4-500 had a G331A mutation outside of the *CalERG11* fragment 4, leading to a D111N amino acid substitution. To check how prevalent this mutation was in the whole library, the *CalERG11* region surrounding the mutation was amplified from the bacterial miniprep for the entire library and sent to Sanger sequencing. The sequencing signal for the wild-type nucleotide at this position was very clear, with no background signal, confirming that this mutation was not present in most F4-500 plasmids.

### Plasmid libraries quality control

The four plasmid libraries were sequenced to check variant diversity. The *CalERG11* F4 from each library was first amplified from 2 ng of each of the bacterial minipreps, with primers adding Illumina index primer binding sites. PCR products were diluted 1:2500 and amplified with Illumina Nextera i5 and i7 primers (Illumina). The PCR reaction was done in triplicate to limit bias in variant frequency introduced by the PCR amplification. Reactions for the same library were pooled and purified with magnetic beads (Axygen AxyPrep Mag PCR Clean-up kit). Equal amounts of each purified sample were then pooled before sending to sequencing. High-throughput sequencing was performed at the Plateforme d’Analyses Génomiques sequencing platform (Institut de biologie intégrative et des systèmes, Université Laval, Canada) using an AVITI system in paired-end 300bp (Element Biosciences). Between ∼281,000-497,000 reads were analysed per library.

Sequencing reads were analyzed using scripts adapted from previous studies (Després et al. 2022; Bédard et al. 2024), and available on the associated Github repository. Python (3.12.9) libraries pandas 2.2.3 (McKinney 2010), numpy 2.5.1 (Harris et al. 2020), matplotlib 3.10.0 (Hunter 2007) and seaborn 0.13.2 (Waskom 2021) were used for data analysis and visualization. The analysis was split into two scripts. The first script was used to evaluate the quality of the demultiplexed reads using FastQC 0.12.1 (Andrews 2010), merge and trim reads according to the primers’ sequence using PANDAseq 2.11 (Masella et al. 2012), aggregate reads for the same sequence using VSEARCH 2.4.4 (Rognes et al. 2016) and align each sequence to the *CalERG11* using Needle from EMBOSS 6.6.0 (Rice et al. 2000). Singleton reads were excluded before the alignment step. The second script was used to find mutations and count reads for each sequence from the aligned sequences files. Unexpected sequences, ie. sequences with a mutation of the first nucleotide or the last two, were discarded, as these positions lie outside of the region that was targeted for random mutagenesis. A vast majority of the sequences present in the four libraries were wild-type (F4-500 : 80.9%, F4-650 : 83.0%, F4-700 : 80.6%, F4-750 : 72.1%). Most sequences were present in low copy number, with a median of 3 reads per unique sequence in F4-700 and F4-750 libraries and 4 reads per sequence for F4-500 and F4-650 libraries.

### Choosing voriconazole concentrations to use for selection on liquid and solid media to screen the random mutagenesis library

A preliminary experiment was performed to choose a voriconazole concentration for selection on solid media. Briefly, we competed two fluorescent *S. cerevisiae* strains, expressing the wild-type copy of *CalERG11* or *CalERG11* with a known resistance-conferring mutation, G465Q (Bédard et al. 2024), in liquid media and plated the culture on solid media containing different voriconazole concentrations to determine which one led to the best enrichment of the resistant variant.

Two fluorescent strains of *S. cerevisiae* were first constructed by tagging *PDC1* in the TetOpr-*ERG11* strain with yeGFP or mCherry (strains TetOpr-*ERG11_PDC1*-yeGFP and TetOpr-*ERG11_PDC1*-mCherry). *PDC1* is highly expressed in yeast, allowing fluorescence to be visualized directly from colonies on agar plates using a UV light transilluminator (Invitrogen SafeImager). Cassettes of yeGFP-HPH and mCherry-HPH were amplified with a forward primer adding homology to the end of the *PDC1* gene and a reverse primer adding homology to the terminator. The yeGFP-HPH cassette was amplified from plasmid pYM25 (Janke et al. 2004) and the mCherry-HPH cassette was amplified from plasmid pBS35 (Ascencio et al. 2021). The amplified cassettes were digested with 20 U of DpnI at 37 °C for 1 h and purified using magnetic beads before transformation in the TetOpr-*ERG11* strain. Transformants were selected on media containing HygB and integration of the cassette was confirmed by performing a yeast colony PCR to amplify the *PDC1*-yeGFP or *PDC1*-mCherry junction and the entire yeGFP or mCherry gene. The TetOpr-*ERG11_PDC1*-mCherry strain was transformed with pRS31N-*CalERG11* (TeOpr-*ERG11_PDC1*-mCherry WT) and the TetOpr-*ERG11_PDC1*-yeGFP strain was transformed with pRS31N-*CalERG11* G465Q (TetOpr-*ERG11_PDC1*-yeGFP G465Q).

Overnight cultures of the two strains were inoculated in 3 mL YPD + Nat. A culture containing 75 % *PDC1*-mCherry WT and 25 % *PDC1*-yeGFP G465Q was prepared and used to inoculate 2 mL YPD + NAT + Dox cultures with voriconazole at 0.08 µg/mL or 0.13 µg/mL at a starting cell density of 0.05 OD. Concentrations of 0.08 and 0.13 µg/mL were chosen as these correspond to the concentrations inhibiting wild-type growth by 50 % (IC50) and 90 % (IC90) respectively, as measured by (Bédard et al. 2024). Cultures were grown to 0.8 OD at 30°C with shaking, then diluted to 0.01 OD. A volume of 100 µL (∼10 000 cells) was plated on YPD + Nat + Dox agar plates with voriconazole at 0.08 µg/mL or 0.4 µg/mL or without voriconazole. Plates were incubated at 30°C for 60 h. TetOpr-*ERG11_PDC1*-mCherry WT and TetOpr-*ERG11_PDC1*-yeGFP G465Q colonies were counted from a picture taken of the plates placed on a transilluminator (Invitrogen SafeImager) to count TetOpr-*ERG11_PDC1*-mCherry WT and TetOpr-*ERG11_PDC1*-yeGFP G465Q colonies. Colonies were counted on one-eighth or one-fourth of the plate. The solid medium concentration had the biggest effect on resistant variant enrichment, as ∼33 % (IC90 liquid) to ∼44 % (IC50 liquid) of colonies were wild-type on plates with 0.08 µg/mL voriconazole while ∼10% (IC90 liquid) to ∼12% (IC50 liquid) were wild-type on plates with 0.4 µg/mL voriconazole. Voriconazole concentrations of 0.08 µg/mL for the liquid culture and 0.4 µg/mL for the solid medium were chosen.

### Screening for resistant *CalERG11* variants in the random mutagenesis libraries

The TetOpr-*ERG11* strain was transformed with the four random mutagenesis plasmid libraries using a standard lithium acetate yeast transformation (Gietz and Schiestl 2007). Transformants were selected using Nat resistance. Around 70,000 colonies were recovered for each library. YPD medium (5 mL) was added to the plates and colonies were scraped using a sterile glass rake. All recovered colonies from the same library were pooled. Aliquots in 25 % glycerol were prepared and stored at -80°C.

All four libraries were grown overnight in 5 mL YPD + Nat media starting with ∼16,000 cells per variant, taking into account that variant sequences represented ∼15-25 % of the sequences in the libraries. Cultures were diluted to 0.05 OD in 5 mL of fresh YPD + Nat + Dox media with 0.08 µg/mL voriconazole and grown to 0.8 OD at 30°C with shaking. Cultures were diluted to 0.05 OD and 100 µL (∼50,000 cells) from each culture were then plated on three YPD + Nat + Dox plates with 0.4 µg/mL voriconazole. Plates were incubated at 30°C for 60 h. The number of colonies was counted on a quarter of each plate to estimate the total number of colonies (F4-500 : ∼4290, F4-650 : ∼3410, F4-700 : ∼3930, F4-750 : ∼2220 total colonies retrieved). A 75 % *PDC1*-mCherry WT and 25 % *PDC1*-yeGFP G465Q mix was grown in the same condition and plated on one YPD + Nat + Dox plate with 0.4 µg/mL voriconazole. *PDC1*-mCherry WT and *PDC1*-yeGFP G465Q colonies were counted on half the plate and it was estimated that *PDC1*-mCherry WT represented ∼7.8% of the colonies.

YPD medium (5 mL) was added to the plates and colonies were scraped using a sterile glass rake. All recovered colonies from the same library were pooled. Aliquots in 25 % glycerol were prepared and stored at -80°C and cell pellets corresponding to an OD of 5 were prepared. Plasmids were extracted using the Zymoprep Yeast Plasmid Miniprep II kit (Zymo Research) following an optimized protocol from (Bédard et al. 2024). The libraries were prepared and sent to sequencing as described in the “Plasmid libraries quality control” section, except the libraries were sequenced with paired-end 275 bp. Between ∼180,000-300,000 reads were analysed per library. Reads processing and variant calling was done as described in the “Plasmid libraries quality control” section. A majority of reads were the wild-type sequence (F4-500 : 70.9%, F4-650 : 74.0%, F4-700 : 66.8%, F4-750 : 75.8%). The median read count per unique sequence was 3 for libraries F4-500 and F4-700 and 4 for libraries F4-650 and F4-750.

### Voriconazole selection data analysis

The variant frequencies after voriconazole selection were calculated using a custom script. The frequency of each sequence was calculated and only sequences found in at least 2 of the 4 libraries were kept. The median frequency across libraries was then calculated for each variant. Mutations leading to the same amino-acid change were counted together only if they were the only mutation present in a given sequence.

Out of the variants retrieved after selection, 326 variants with a single mutation had previously been characterized by (Bédard et al. 2024) and classified as beneficial, neutral or deleterious based on a comparison of the variant’s fitness to the wild-type in a competition experiment with voriconazole. Here, we classified the beneficial variants as “resistant” and neutral or deleterious variants as “sensitive”. We used the distribution of the frequencies after selection of the known sensitive variants to set a threshold value above which resistant variants were enriched, in order to facilitate identification of potential resistance-conferring mutations to validate. The threshold was set at the 99th percentile of the frequencies of sensitive variants.

To compare the variant frequencies before and after selection, the frequency of the variants before the selection was calculated as the median frequency of the translated amino-acid sequence in all the plasmid libraries in which the variant was sequenced. We excluded 550 variants, including seven known variants, from this comparison, as they were found after the selection but were not sequenced in the initial plasmid libraries. Variant frequencies before and after selection were highly correlated (Spearman’s rank correlation, ρ = 0.62, p = 1.31 x 10^-115^). This is also the case for the wild-type sequence, which represents a majority of the sequencing reads before and after selection.

### Validation of variants identified in the voriconazole selection

Nine variants were individually reconstructed by site-directed mutagenesis as described in the “Cloning” section. The presence of the mutation was confirmed with Sanger sequencing. Growth in liquid YPD media containing Nat, Dox and voriconazole (0.086 µg/mL) was measured as described in the “Growth curves” section, with five replicates per variant. Voriconazole concentration represents the IC50 of the wild-type, calculated following (Bédard et al. 2024), for the new batch of voriconazole that had to be prepared for this experiment. Growth curves data was analyzed using a different custom script as the one described in the “Growth curves” section, adapted from (Bédard et al. 2024). OD measures were corrected for variation in initial cell density by subtracting the median of the first 3 measurements. AUC was calculated for the first 30 h of growth and normalized by the average AUC of the wild-type. Growth of the variants was compared to the wild-type using a two-sided independent t-test with correction for multiple comparisons using the Benjamini-Hochberg method. Variants with a corrected p-value < 0.05 and AUC > 1 were considered resistant. Variants with lower (corrected p-value < 0.05 and AUC < 1) or similar (corrected p-value > 0.05) growth compared to the wild-type were considered sensitive.

### Protein structure visualization

The crystal structure of *Cal*Erg11p bound to itraconazole was taken from the Protein Data Bank (PDB ID : 5V5Z, (Keniya et al. 2018)). ChimeraX v1.12 (Meng et al. 2023) was used to visualize and annotate the protein structure. For residues R414, Y415, K451 and P515, all possible atom-to-atom distances with the itraconazole and the heme molecules were calculated using the “contacts” command with “distOnly” and “log True” options. The shortest distances between the residue and both molecules were manually retrieved from the ChimeraX log

## Supplementary Figures

**Supplementary Figure 1.**
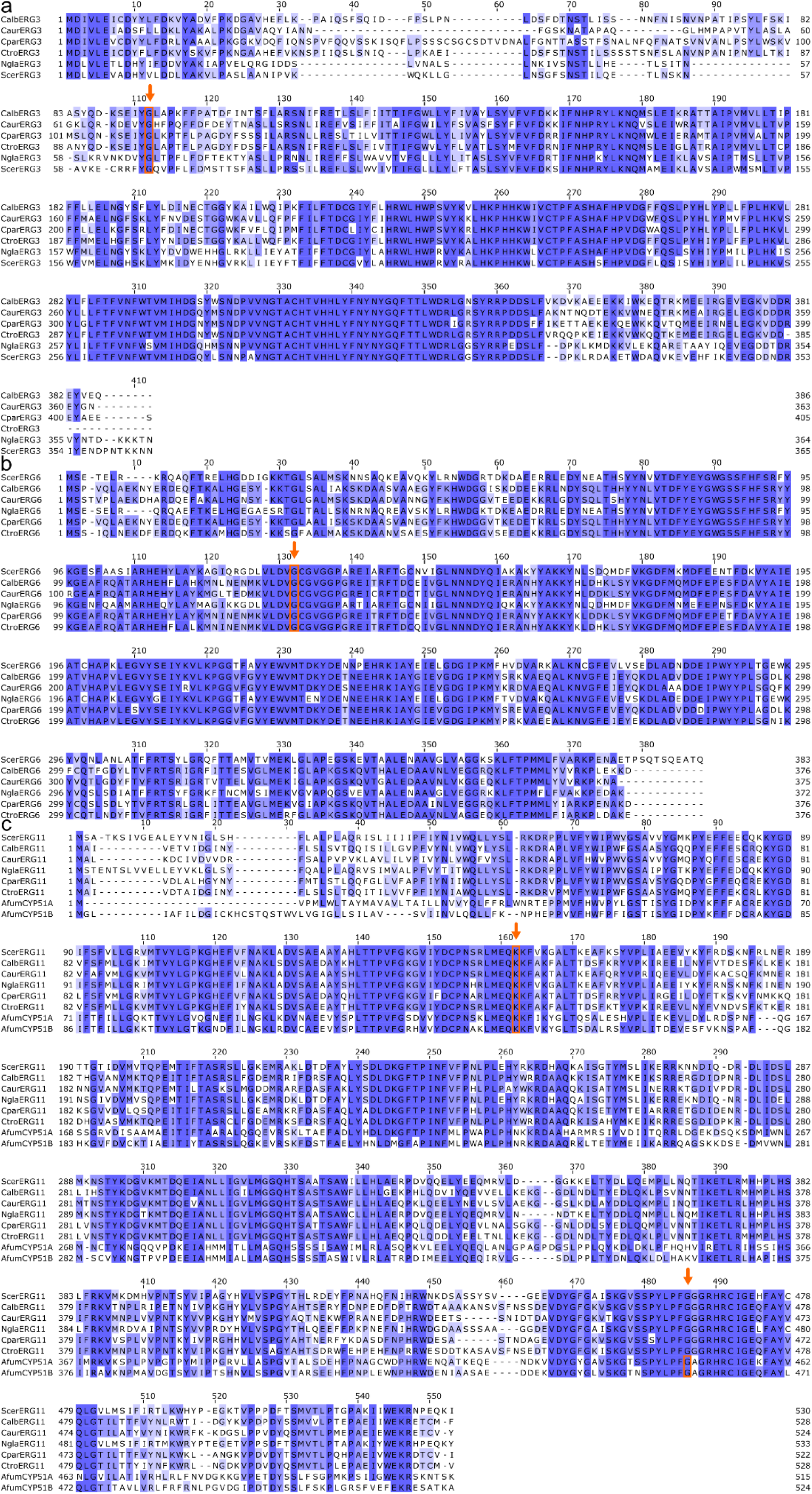
Multiple sequence alignments of protein sequences of orthologs used in this study, composed of a) Erg3p sequences, b) Erg6p sequences and c) Erg11p sequences. Colored by conservation, with darker blue indicating greater conservation of the residue at a given position. Positions in which a mutation was built are indicated with orange boxes and arrows.

**Supplementary Figure 2.**
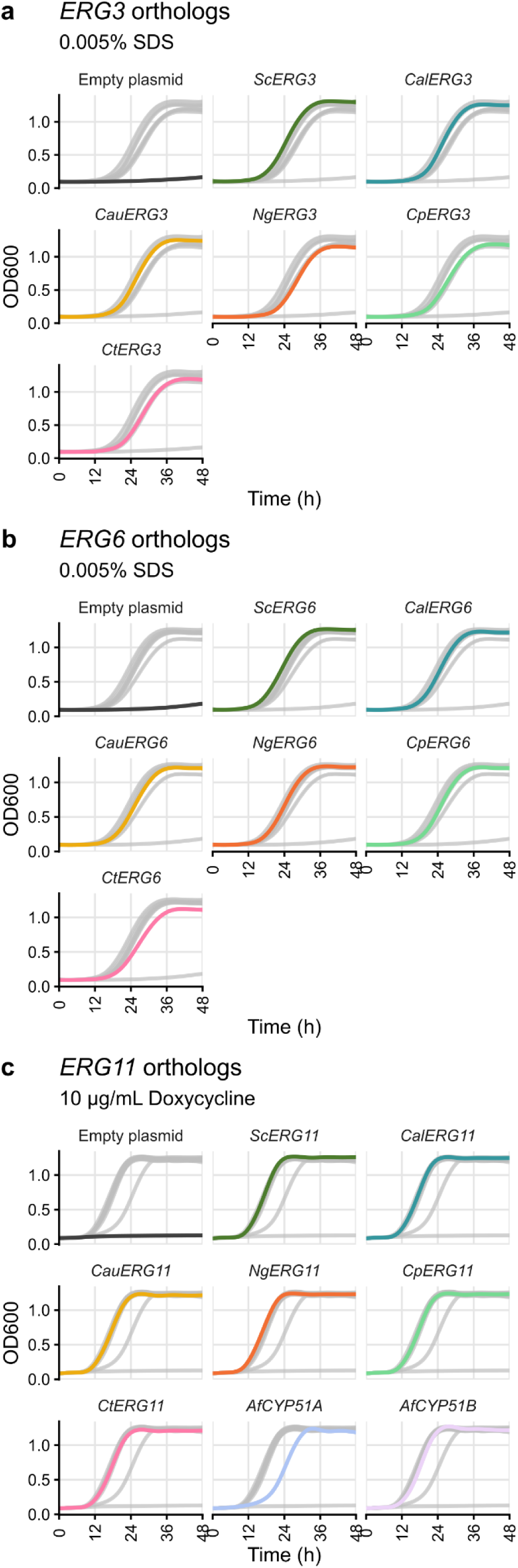
Raw growth curves measuring the complementation of the *S. cerevisiae* deletion or repression by heterologous expression of orthologs of a) *ERG3* in the *erg3Δ*::*HPHNT1* strain, b) *ERG6* in the *erg6Δ*::*HPHNT1* strain, and c) *ERG11* in the TetOpr-*ERG11* strain. Each of the subplots shows one ortholog (or the empty plasmid control) in color, while all the other orthologs are shown in the background in light grey.

**Supplementary Figure 3.**
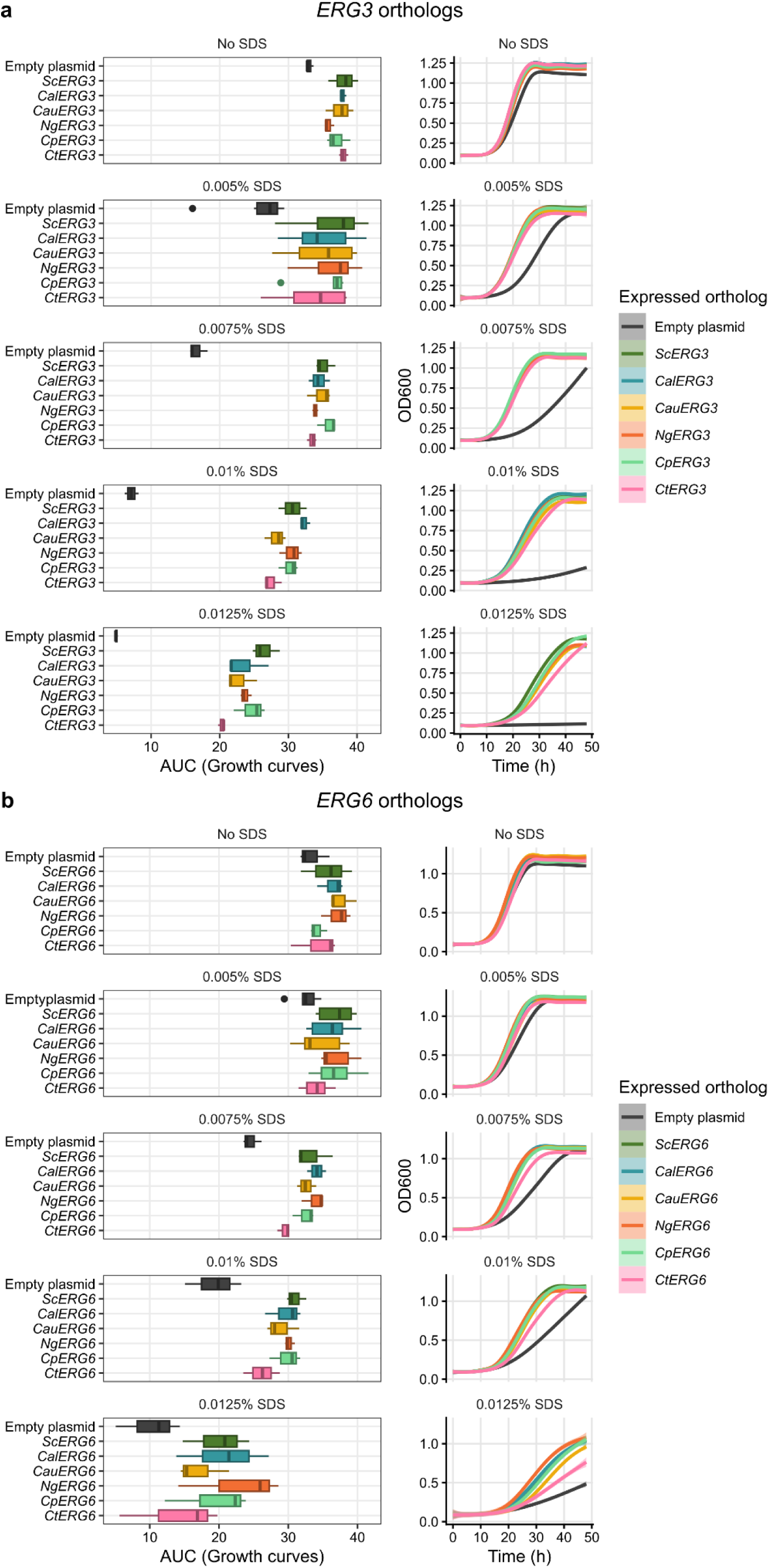
Increasing the concentration of SDS leads to clearer complementation by TetOpr-*ERG3* and TetOpr-*ERG6* Area under the curve (AUC) measurements (left) and the raw growth curves from which they were calculated (right) of the a) TetOpr-*ERG3* and b) TetOpr-*ERG6* strains containing plasmids of the heterologous expression system, at different concentrations of SDS. AUCs and growth curves calculated from 3 replicates.

**Supplementary Figure 4.**
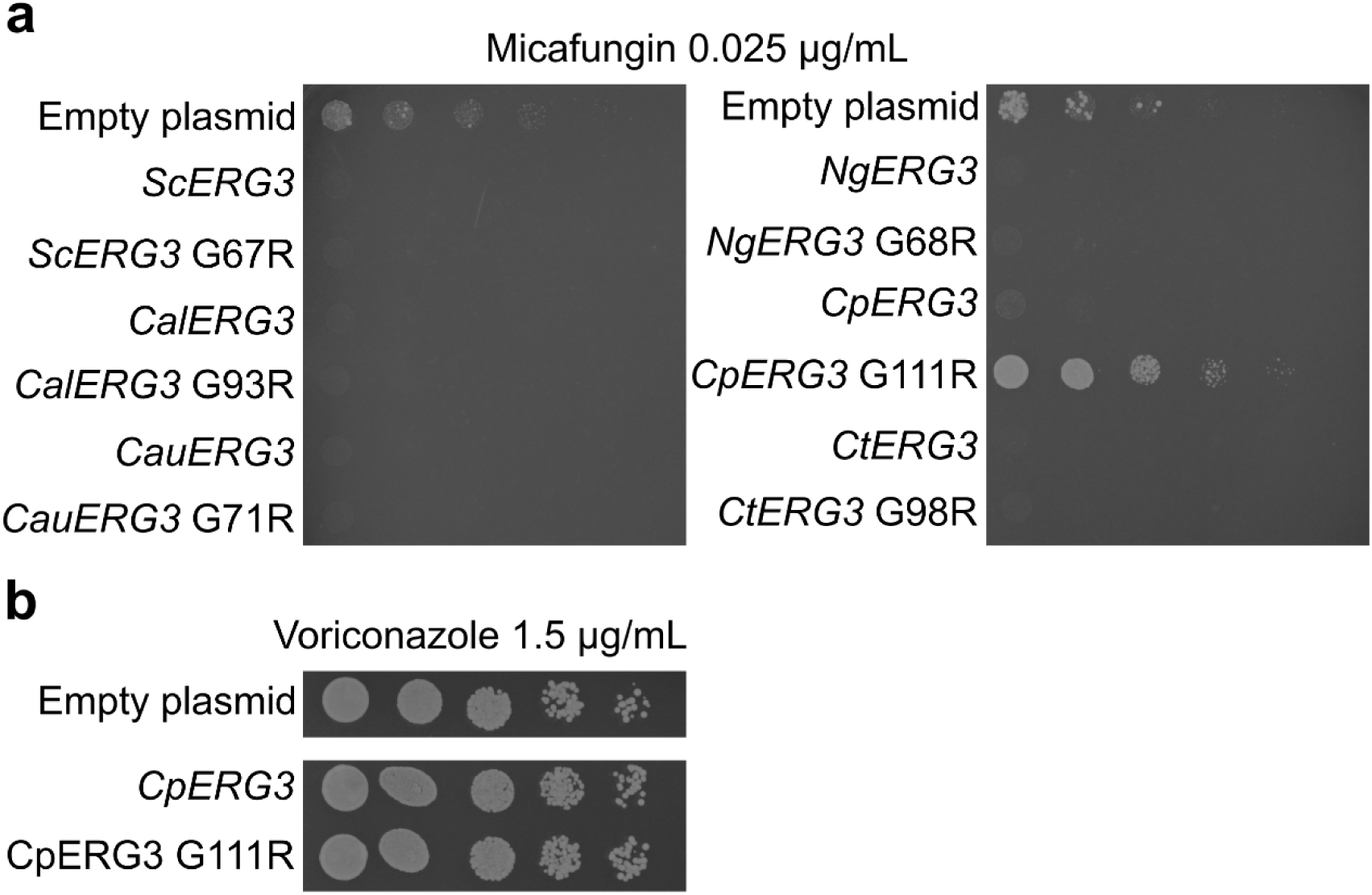
*CpERG3* G111R is the only tested mutant resistant to micafungin, but is not more resistant than the wild-type to voriconazole. Spot dilution assays showing susceptibility, or lack thereof, of empty plasmids, wild-type and mutant orthologs of a) multiple orthologs of *ERG3* to 0.025 µg/mL of micafungin, b) *CpERG3* to 1,5 µg/mL of voriconazole

**Supplementary Figure 5.**
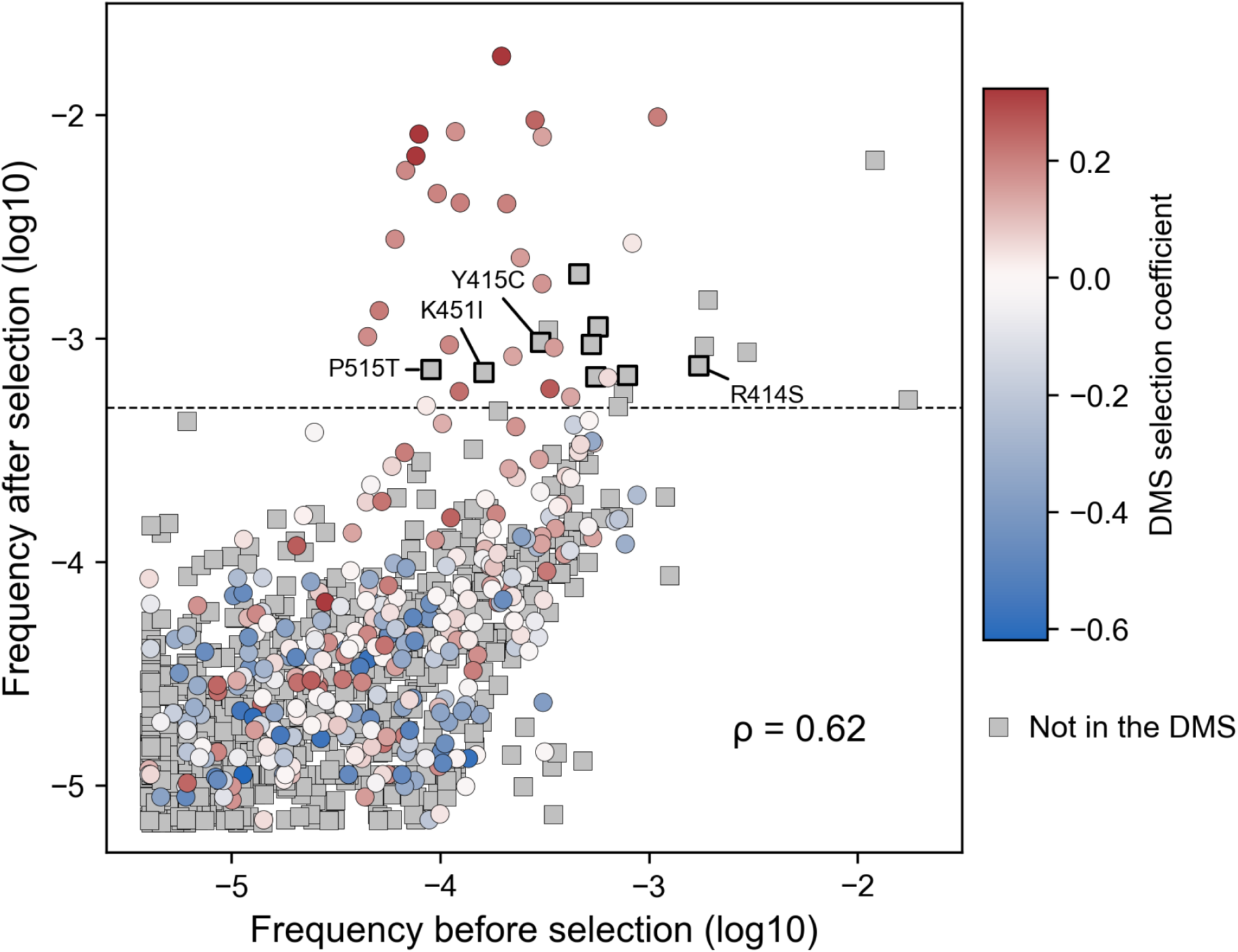
Relationship between the variant frequencies generated by random mutagenesis before and after voriconazole selection. Variant frequency before selection is the median frequency of a variant for all plasmid libraries in which it was sequenced and of synonymous codons for a given amino acid substitution, when available. Variant frequency after selection is the median of two to four selection experiments and of synonymous codons for a given amino acid substitution when available. Variants previously characterized by deep mutational scanning (DMS) in (Bédard et al. 2024) are represented by circles and colored according to their selection coefficients as calculated in that study, with positive values representing higher fitness than the wild-type in a competition experiment with voriconazole and negative values representing lower fitness. Variants not previously characterized by DMS are represented by grey squares. Black dotted line represents the threshold value used to select variants to validate. Variants selected for further characterization (Figure 5d) have a thicker black outline and mutations validated to confer resistance are identified. Two-sided Spearman’s rank correlation is shown (p = 1.31 x 10^-115^).

## Acknowledgments

The authors would like to thank members of the Landry research group for valuable comments and feedback. In particular, we thank Mégane Bernier for construction of the pRS31N-*AfCYP51B* plasmid.

## Data availability

Strains are available upon request. All plasmids will be submitted to Addgene for the final submission of this article. All scripts used and described in this paper, and all scripts used to make plots are available in the associated GitHub repository (https://github.com/Landrylab/Jordan-et-al-2026). Liquid growth data (growth curve) and protein sequence alignments can also be found in the associated Github repository (https://github.com/Landrylab/Jordan-et-al-2026). Sequencing reads from the random mutagenesis library, and from the mutants after selection by voriconazole can be found on the SRA (PRJNA1527525). All figures, supplementary figures, raw as well as cropped and grayscale images used in making figures can be found on figshare (https://figshare.com/s/179c9805f571e84f1963). Table S1 contains details of all growth media used. Table S2 contains details of all protein and gene sequences used for the *ERG3, ERG6* and *ERG11* orthologs. Table S3 contains details of all strains, plasmids and primers used.

